# *In vitro* impact of fluconazole on oral microbial communities, bacterial growth and biofilm formation

**DOI:** 10.1101/2023.08.03.551749

**Authors:** Louise Morais Dornelas-Figueira, Antônio Pedro Ricomini Filho, Roger Junges, Heidi Aarø Åmdal, Altair Antoninha Del Bel Cury, Fernanda Cristina Petersen

## Abstract

Anti-fungal agents are widely used to specifically eliminate infections by fungal pathogens. However, the specificity of anti-fungal agents has been challenged by a few studies demonstrating anti-bacterial inhibitory effects against Mycobateria and Streptomyces species. Here we evaluated for the first time the potential effect of fluconazole, the most clinically used antifungal agent, on a human oral microbiota biofilm model. The results showed that biofilm viability on blood and mitis-salivarius agar media was progressively increased in the presence of fluconazole at clinically relevant concentrations, despite of a reduction in biomass. Target PCR revealed higher proportions of *Veillonella atypica, Veillonella dispar* and *Lactobacillus* spp. in the fluconazole treated samples compared to the control, while *Fusobacterium nucleatum* was reduced and *Streptococcus* spp was not significantly affected. Further, we tested the potential impact of fluconazole using single-species models. Our results using *Streptococcus mutans* and *Streptococcus mitis* luciferase reporters showed that *S. mutans* planktonic growth was not significantly affected by fluconazole, while for *S, mitis* planktonic growth, but not biofilm viability, was inhibited at the highest concentration. Fluconazole effects on *S. mitis* biofilm biomass were concentration and time-dependent. Exposure for 48h to the highest concentration of fluconazole was associated with *S. mitis* biofilms with the most increased biomass. Potential growth inhibitory effects were further tested using four non-streptococcal species. Among these, planktonic growth of both *Escherichia coli* and *Granulicatella adiacens* were inhibited by fluconazole. Conclusions: The data indicate bacterial responses to fluconazole that extend to a broader range of bacterial species than previously anticipated from the literature, with the potential to disturb microbial biofilm communities. Future studies are warranted to further identify the breath of species for which growth may be impacted by anti-fungal agents, and possible mechanisms involved.

## 1 Introduction

The oral cavity is colonized by numerous microorganisms, including bacteria, fungi, archaea, protozoa, and viruses. Together, they form what is known as the oral microbiota. The coexistence of these microorganisms relies on their synergistic and antagonist interactions, resulting in a balanced microbial community that maintains potentially harmful microorganisms numbers at low levels(1). However, stressors can disturb this microbial balance, leading to a condition called dysbiosis(2). Dysbiosis results in an altered microbial community where potentially pathogenic microorganisms become more prevalent, increasing the risk of disease. Besides, commensals play an important role in the establishment and balance of the microbiota and, once this dynamic balance is disrupted, infections take place (3). One of the most important stressors affecting microbial communities is the use of antimicrobials agents (1,4).

Antimicrobials play a crucial role in the treatment and control of infectious diseases. This includes antifungal agents such as fluconazole, one of the most effective and prescribed agents for antifungal prophylaxis and for the treatment of oropharyngeal candidiasis in HIV-positive patients, neonatal and *Candida* spp. infections (5–10). Fluconazole has been used successfully to treat local diseases, such as vaginal candidiasis, oropharyngeal and esophageal candidiasis, and *Candida* spp. urinary tract infections; but also, to treat systemic infections including candidemia, disseminated candidiasis, coccidioidomycosis and cryptococcal meningitis (6,11,12).

Fluconazole is a triazole that inhibits the cytochrome P450 enzyme lanosterol demethylase (14α-demethylase) involved in the ergosterol biosynthesis pathway in fungi, thus inhibiting cell membrane formation (5). It is available mostly as enteral and intravenous preparations, but also as mouthrinse or suspension for local infections (13,14). The clinical efficacy of systemic fluconazole in preventing and treating oropharyngeal and esophageal candidiasis is attributed to the relatively high concentrations achieved in salivary secretions following oral administration (15).

The impact of antimicrobials in the human microbiota is an area that is receiving increasing attention due to the association of microbial community imbalances with different diseases and the alarming levels of antimicrobial resistance. Antifungals have the potential to disturb the balance of microbial communities in the gut (16) and promote diarrhea and nausea. In animal models, disturbance of fungal populations leading to gut microbiota dysbiosis has been associated with induction of host pro-inflammatory responses (17). Also, increasing studies indicate that certain antifungal drugs, in particular imidazoles, may have direct antimicrobial activity against selected bacterial pathogens and commensals of the gut (16). For antifungal triazoles, including fluconazole, antimicrobial activity against gut microbes has not been observed using stringent cutoffs (16). It is possible, however, that fluconazole may have weak inhibitory activity, as observed for *Streptomyces lividans*, a soil bacterium also found in the gut (18). Such studies indicate that antifungals may potentially promote dysbiosis in complex microbial communities by a third and unexplored mechanism, namely through antibacterial effects.

In contrast to the gut, negative effects of antimicrobials on the microbiota of the upper digestive tract are less understood. This is a field of increased interest due to the association of oral microbiome dysbiosis with various autoimmune, inflammatory, and neoplastic conditions (19–21). Also, the microbial community of the upper digestive tract represent a reservoir of antibiotic resistance genes and bacterial pathogens (22–24). In this study we aimed to investigate the possible effect of fluconazole (A) on a complex human microbial community using an oral microbiota biofilm model, and (B) on growth of selected species of streptococcal and non-streptococcal colonizers of the oral cavity.

## 2 Materials & Methods

### 2.1 Experimental Design

For the oral microbiota study, non-stimulated saliva were collected from 6 healthy subjects aged between 25-35 years old. Subjects were asked to abstain from drinking and eating 2 hours prior to saliva collection. The study was conducted in accordance with the Declaration of Helsinki and approved by the Norwegian Regional Ethics Committee (REK20152491) for studies involving human samples. The saliva were first collected, centrifuged, pooled and added to the bottom of the wells of 24-well plates to form a salivary pellicle, as previously described (25). After sterilization in UV-light (2500 μJ/cm^2^ for 30 min) SHI media containing pooled salivary microbiota (2 μl/ml) was added to the wells and incubated in an anaerobic chamber at 37°C for 24 h. The detached cells and old media were then carefully removed, and fresh SHI medium containing fluconazole (Cayman Chemical Company; Michigan, USA. 11594) was added to the wells. Fluconazole stock solutions prepared by suspending 5 mg fluconazole in 5 mL 100% ethanol were stored in -20°C. Two concentrations of fluconazole were tested in the microbiota model: 2.56 μg/mL (concentration in saliva after oral administration (SA); Force and Nahata, 1995) and 2000 μg/mL (concentration in mouthrinses - MR) (13). Samples without fluconazole, but with the same concentration of the dissolving agent, were included as negative controls. Three independent experiments, with two to three replicates were performed. After 24h incubation, the biofilm supernatants were analyzed for potential effects of fluconazole on metabolic activities by measuring final pH. Biofilms were resuspended in PBS and used for (1) measurement of potential effects on microbial viability by determining colony forming units (CFU) on agar plates after 48 h incubation at 37°C on 5% CO_2_ atmosphere, (2) measurement of bacterial and fungal DNA by quantitative PCR, (3) measurement of changes in microbial composition by targeted PCR and (4) biofilm dry biomass by weight measurements.

To evaluate the effects of fluconazole on *Streptococcus* spp. planktonic growth, we used the type of strain of *S. mitis* and the *S. mutans* strain UA159, each engineered to harbor an ldh-luciferase reporter system, as described below. The impact of fluconazole on *S. mitis* biofilm growth was also investigated. In this assay, biofilms were formed in 96-well flat bottom plates with fluconazole at different concentrations: saliva concentration 2.56 μg/mL (SA), peak plasma concentration 4.39 μg/mL (1X), twofold peak plasma concentration 8.78 μg/mL (2x) and at mouthrinse concentration 2.0 mg/mL (MO).The microorganisms were incubated at 37°C and the relative light units (RLU) and optical density (OD) were measured every 30 min. Negative control groups without fluconazole were included in all experiments. Three independent experiments, with two to three replicates were performed. To elucidate the effect of fluconazole on *S. mitis* biofilm formation, biofilms were grown in tryptic soy broth (TSB) supplemented with 0.2% glucose in 5% CO_2_ at 37°C. After 6, 24 and 48 h the biofilms were harvested and CFU, biomass (dry weight) and pH were measured.

### 2.2 pH assay

The pH in the supernatant of the oral microbiota samples was measured at 6, 24 and 48 h growth with the aid of a pH meter (pH 3110 Einzelgerät - WTW®) calibrated with pH standards of 4.0 and 7.0.

### 2.3 Viability Assay

To estimate the viability of microorganisms in the oral microbiome samples, the counts of the colony forming unit (CFU) were performed. Aliquots of 100 μL of the biofilm suspension were ten-fold serially diluted in PBS until 10^−7^. Two separate drops of 20 μL of the dilution were plated on both Blood Agar and Mitis Salivarius Agar (MSA) and incubated at 37°C, 5% CO_2_ for 24 h. The count was carried out with the aid of a stereoscopic microscope.

### 2.4 Biomass

Dry weight was used to evaluate the potential impact of fluconazole on oral microbiome biomass. An aliquot of the suspension was transferred to previously weighed microtubes. Later an amount of 3 times absolute ethanol was added to the tubes and stored in -20°C for 20 min. Then, the tubes were centrifuged at 10.000 *g* for 5 min at 4°C, the supernatants were discarded, and the biofilms were centrifuged under heat and vacuum to dehydrate the samples. Dry weight was determined by the difference of the final and initial microtube weight.

### 2.5 Real-time PCR

To evaluate the concentration of bacterial and fungal DNA in the microbiota, total DNA was extracted using the Zymo Quick-DNA Fungal/Bacterial Micro prep extraction kit (Zymo Research) and quantified by real-time PCR using Zymo DNA quantification kits for bacteria/fungi (Femto Bacterial and fungal DNA Quantification kit, Zymo Research) following the manufacturer’s protocol. The samples used for DNA extraction were from 200 μL of the 1mL re-suspended biofilms. For the PCR reactions, we used 3μL of undiluted DNA for measuring the amount of fungal DNA and 3 μL of 1000-fold dilutions for measurement of bacterial DNA, added to 18 μl of the Femto™ Fungal/Bacterial qPCR Premix. “No template” controls were also included. The AriaMx Real-Time PCR software system was used for calculations (Agilent Techonologies).

Real-time PCR was further used to investigate the impact of fluconazole on microbial composition, using primers specific to *Streptococcus* spp., *Lactobacillus* spp., *Veillonella atypica, Veillonella dispar, Prevotella intermedia* and *Fusobacterium nucleatum* (Table). The samples used were the same as above, but with the DNA adjusted to have similar concentrations in the reactions. The final PCR reaction volume was 25 μl, comprising 12.5 μl Maxima SYBR Green/ROX qPCR Master Mix (2×) containing Maxima Hot Start Taq DNA Polymerase, dNTPs and SYBR Green I in an optimized PCR buffer with ROX passive reference dye, 3 ng DNA template, 0.4 μM forward and reverse primers, The thermal cycling program was as follows: 95°C for 10 min; and then 40 cycles consisting of denaturation at 95°C for 15 s, primer annealing at 55°C for 30 s, and primer extension at 72°C for 30 s. The thermal cycle was finalized with a cycle of 95°C for 1 min, 55°C for 30 s, and 95°C for 30 s. Dissociation curves were prepared immediately after the last PCR cycle. Relative fold changes were calculated using the 2–ΔΔCt method (26).

### 2.6 Construction of luciferase reporters

Lactate dehydrogenase is essential in the ATP generating pathway of streptococci. Its promoter activity provides both viability and metabolic status information in response to antimicrobials (27). The luciferase reporter in *S. mutans* (SM120; table 1) was constructed using the pair of primers FP514 and FP515 to amplify the lactate dehydrogenase promoter region (p*ldh*) of *S. mutans* UA159 (**Table 1**). The primers were designed with the *Nhe*I and *Bam*HI restriction sites on the 5’ end. The resulting amplicon was digested with the referred restriction enzymes and ligated to pFW5-luc (28), a luciferase reporter plasmid carrying a spectinomycin resistance cassette that works in both Gram+ and Gram-bacteria. Cloning in *E. coli* was performed as described previously (29), and a plasmid with the correct insert was then purified and used to transform *S. mutans* UA159 via natural transformation (30). Insertion in the *S. mutans* chromosome was confirmed by selection of positive mutant colonies performed in tryptic soy agar plates with spectinomycin (500 μg/ml) and verified phenotypically using the luciferase growth assay. Design and construction of mutant MI048 in *S. mitis* has been previously described (31).

**Table 1.**
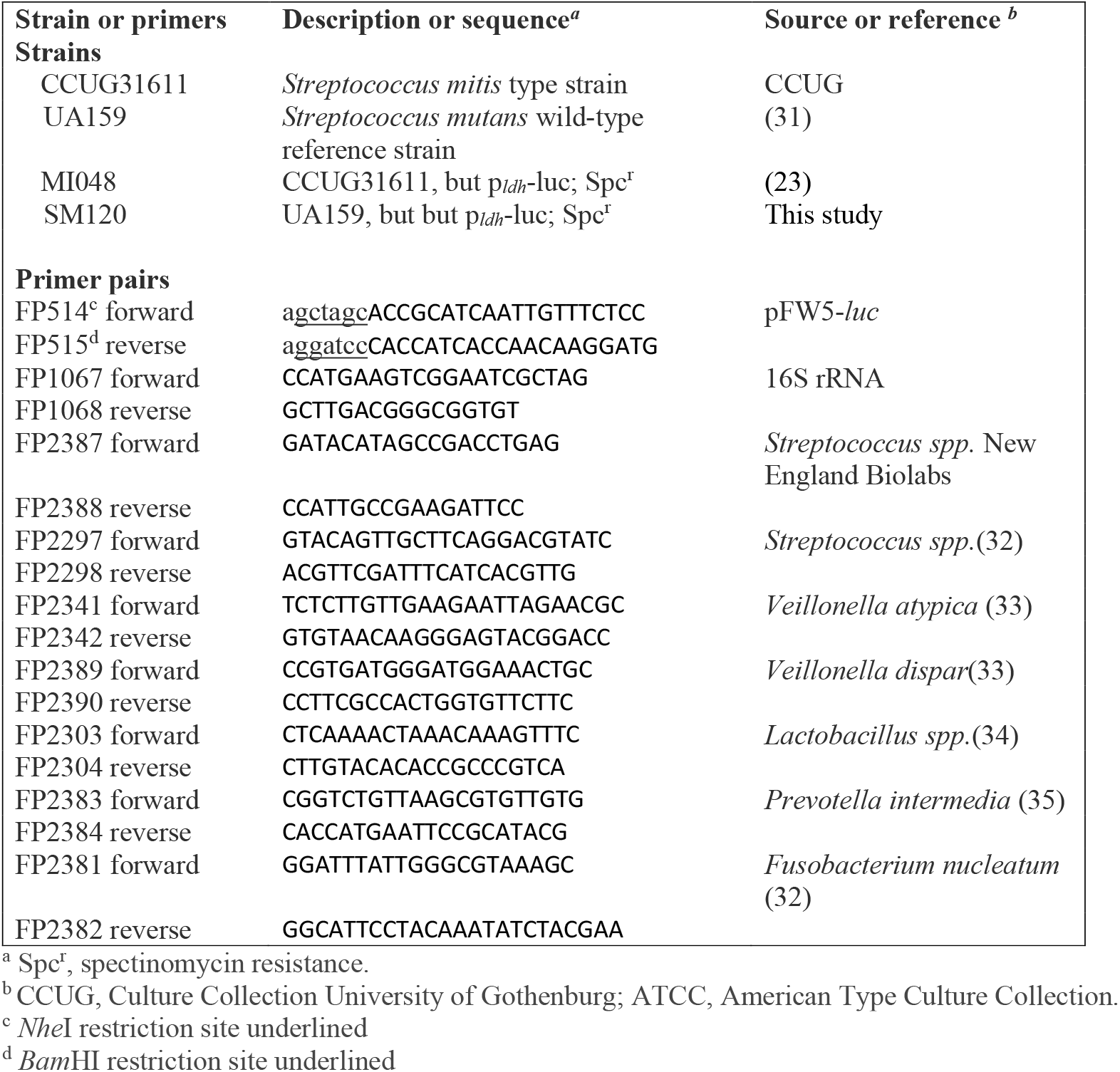
Bacterial strains and primers used in this study.

### 2.7 Luciferase Reporter Assay

The luciferase reporter assay was as described before (31). Briefly, overnight cultures of *S. mitis* MI048 and *S. mutans* SM120 reporter strains were adjusted to the final concentration of approximately 1 × 10^8^ mL^-1^ (OD_600_ 0.1) (31). The microorganisms were cultured in 96-well flat bottom plates (Nunc Thermo Scientific) with tryptic soy broth (TSB) supplemented with 0,2% glucose and different fluconazole concentrations. A negative control group without fluconazole was also included, as well as blanks containing pure medium. In addition, 10 μl room temperature 1.0 mM luciferin solution (Synchem, Felsberg-Altenberg, Germany) was added to each 200 μl culture. The cells were incubated at 37°C and the relative light unit (RLU) and optical density (OD) were measured at various time intervals during growth in a microplate reader (Synergy HT; BioTek, Winooski, VT).

### 2.8 Planktonic growth

To better elucidate whether fluconazole effect on bacterial growth was restricted to *Streptococcus* spp., a planktonic growth assay was conducted with Escherichia coli (DH10BTM), *Granulicatella adiacens* (clinical isolate), *Lactobacillus crispatus* (ATCC 33820) and Lactobacillus salivarius (ATCC 11741). Overnight cultures were diluted to OD600 0.1 and were incubated at 37°C until they reached stationary phase. Optical density (OD) was measured at various time intervals in a Bioscreen C system reader (Lab systems Helsinki, Finland).

### 2.9 *S. mitis* Biofilm Assay

We further analyzed the effects of fluconazole on biofilm formation by *Streptococcus mitis*. For that, we used the *S. mitis* strain used in the planktonic experiments described above. Frozen stock cultures were used for growth in blood agar plates. After 24 h, colonies were transferred to tryptic soy broth (TSB) supplemented with 0.2% glucose and incubated in 5% CO_2_ at 37°C. The inoculum was adjusted to the final concentration of OD 0.1 and the biofilms were formed in 96-well flat bottom plates with fluconazole, using the same concentrations as for planktonic growth. The biofilms were assessed for bacterial viability (CFU counts) and biomass (dry weight) at 6, 24, and 48 h.

### 2.10 Statistical analyses

The results were analyzed using GraphPad software. The assumptions of equality of variances and normal distribution of errors were evaluated for each variable. When the data was not normally distributed, they were transformed. The statistical analysis was performed using one-way ANOVA, having fluconazole concentration as study factor. Post-ANOVA comparisons were performed using the Tukey test.

## 3 Results

### 3.1 Oral Microbiota Biofilm

The evaluation of the fluconazole effects on oral microbiota showed that the drug had no significant effect on the pH of biofilm supernatants collected at 24h after treatment, despite a tendency for lower pH in the absence of fluconazole (p>0.05) (Figure 1A). In contrast, the highest concentration of fluconazole (2000 μg.mL^-1^) had a significant effect on biofilm biomass (p<0.05) (Figure 1B). Regarding the CFU in the biofilms, the mouthrinse concentration (2000 μg.mL^-1^) resulted in increased viable counts on both blood agar and Mitis salivarius agar compared to the control, indicating an effect of fluconazole on the overall microbial community viability (blood agar plates), including *Streptococcus* spp (Mitis salivarius agar plates) (p<0.05) (**Figure 1C,D**).

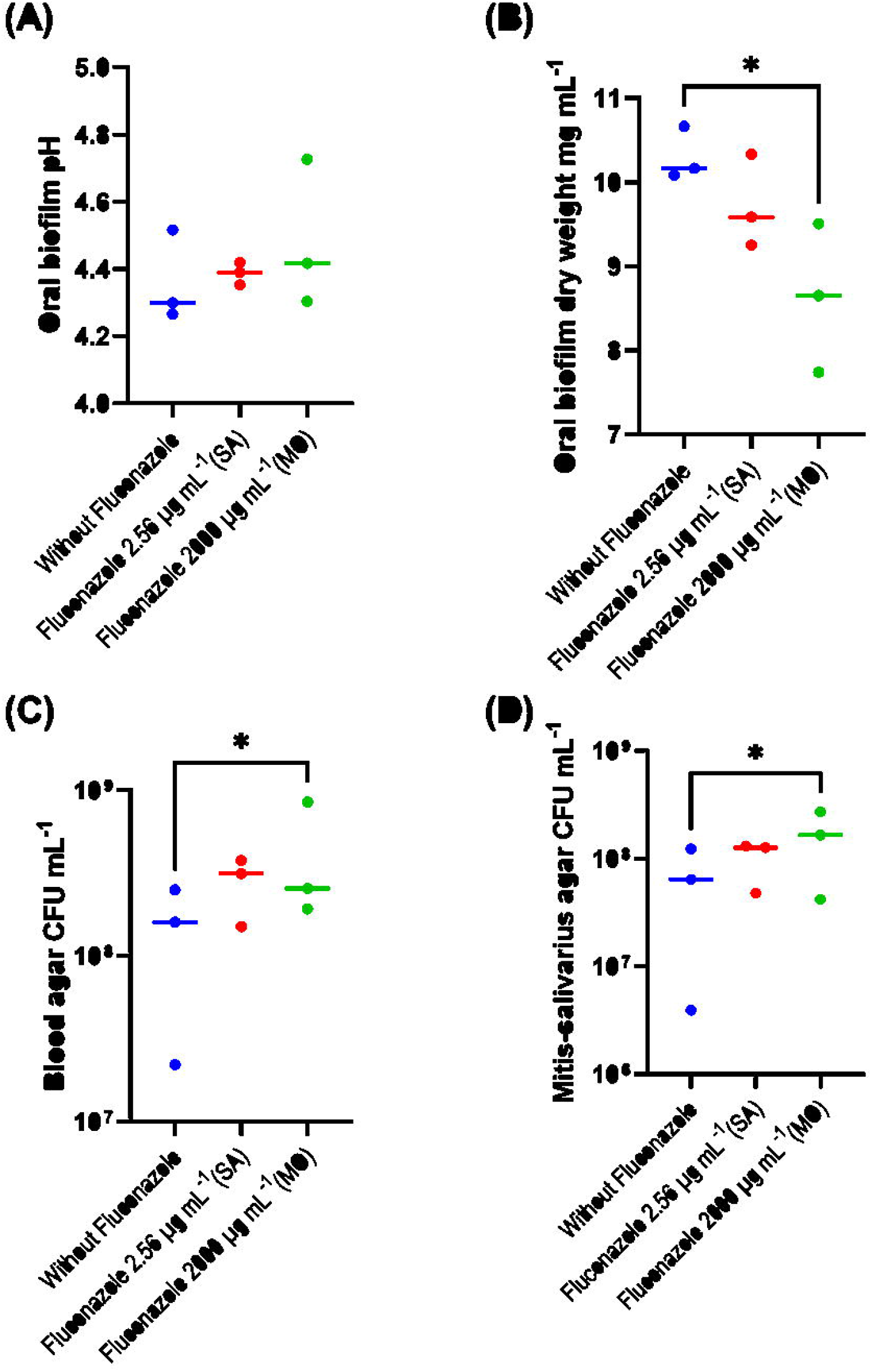

qPCR results with quantitative sensitivity down to 20 femtogram did not reveal significant effects of fluconazole on total bacterial DNA (Figure 2). For fungal DNA, the amounts measured were below 20 fg demonstrating that fungal DNA was more than nine orders of magnitude lower than the bacterial DNA in the samples. For the target PCR using three biological replicates for each group, only *Prevotella intermedia* was below detection level. The largest changes were for *V. atypica, V. dispar* and *Lactobacillus*, which were present in higher amounts in the fluconazole treated samples compared to the control, while *F. nucleatum* was reduced in the treated group (Figure 3).

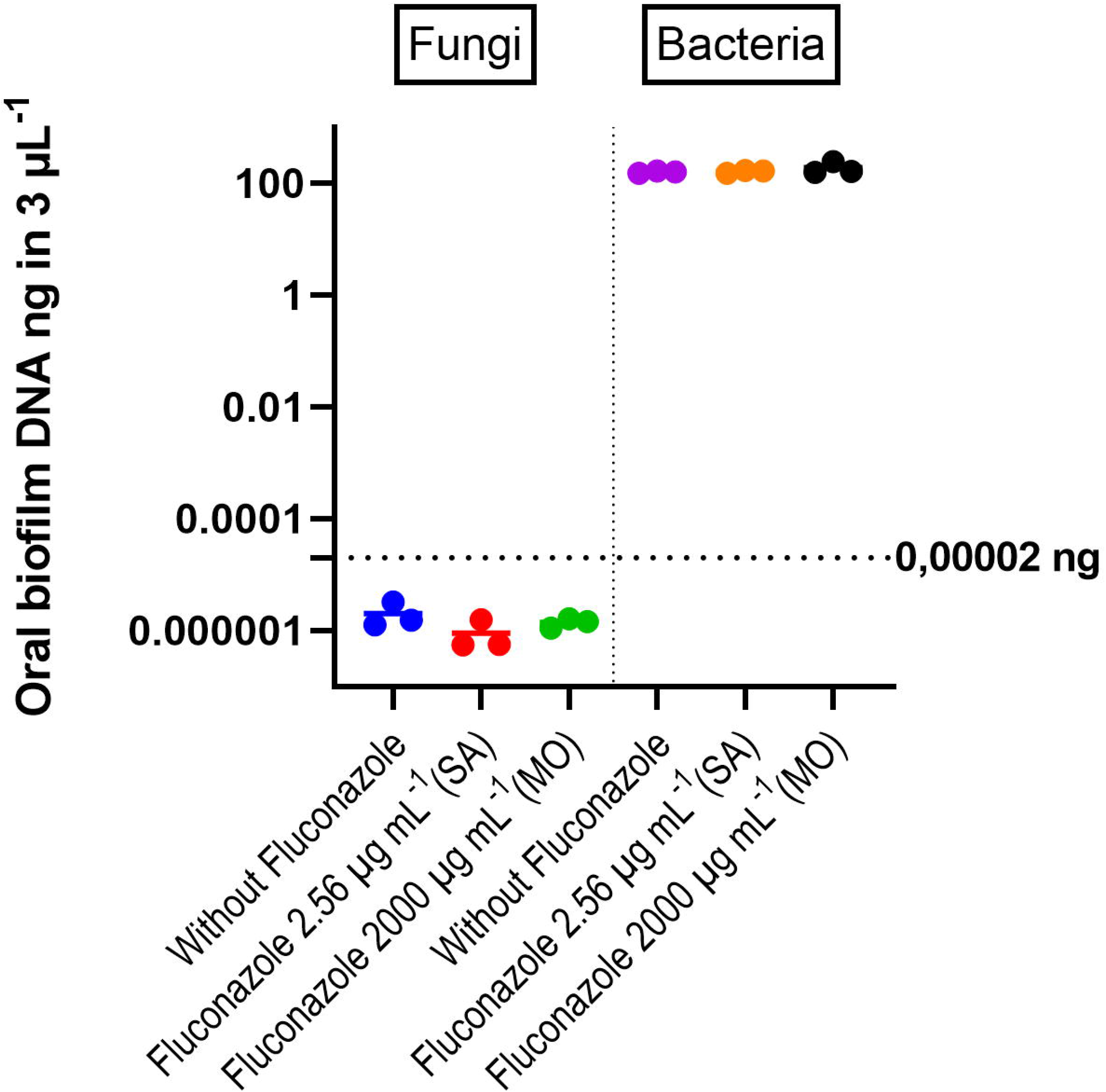

**Figure 3:**
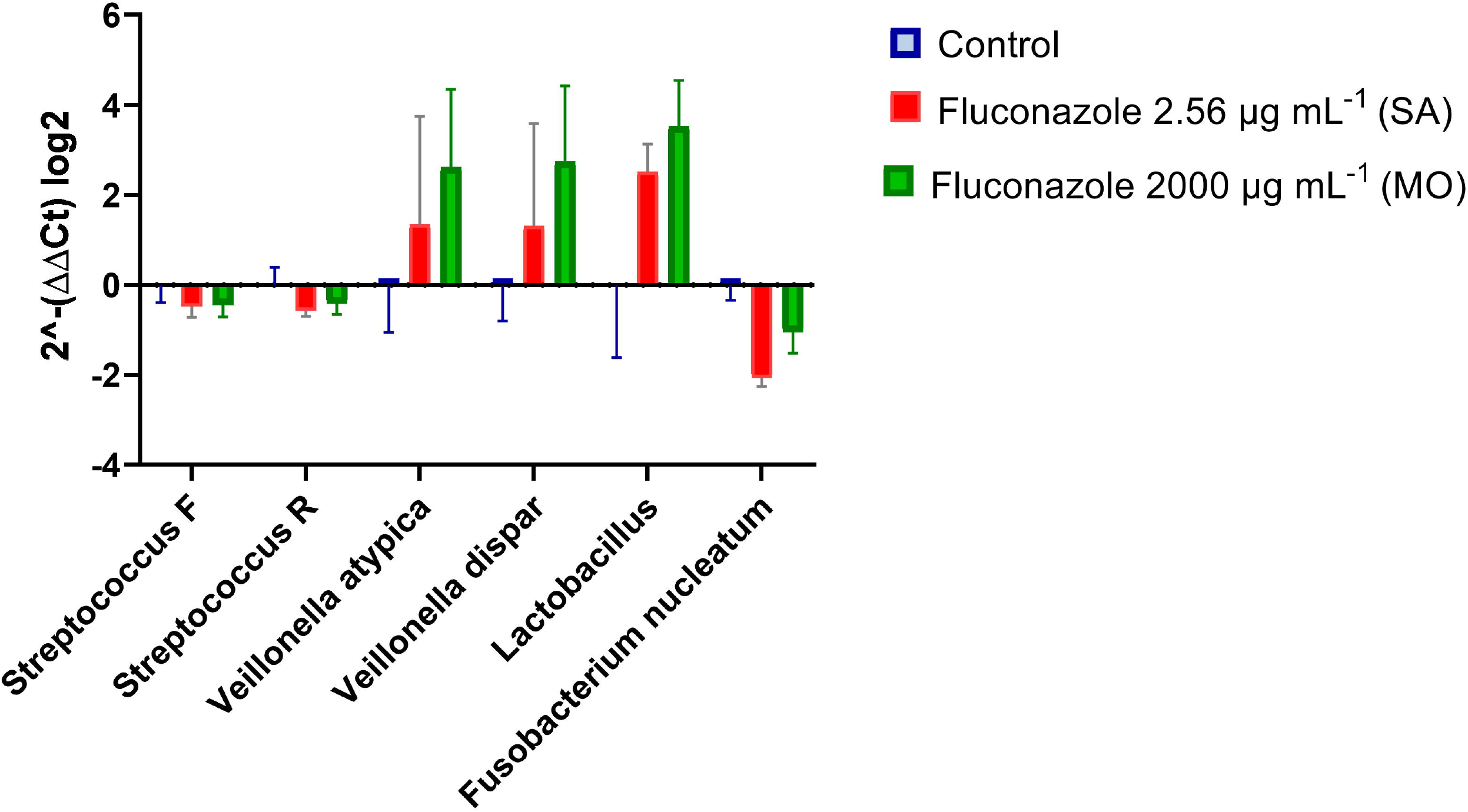
Real-time PCR results expressed by relative fold changes (2^−ΔΔCt^) show the impact of fluconazole at 2.56 μg.mL^-1^ (SA), 2000 μg.mL^-1^ (MO), or without fluconazole exposure (negative control) on microbial composition. *Veillonella atypica, Veillonella dispar and Lactobacillus* spp., were present in higher amounts in the fluconazole treated samples compared to the control, while *F. nucleatum* was reduced, indicating that fluconazole changes microbiome composition.

### 3.2 Fluconazole effect on bacterial planktonic growth

The luciferase assay revealed a dose-dependent reduction on viability and metabolic activity of *S. mitis* in the presence of fluconazole **(Figure 4A)**. Growth measured as density was also inhibited **(Figure 4B)**. For *S. mutans*, no significant effects were observed (**Figure 4C and 4D**). Interestingly, fluconazole reduced planktonic growth of *E. coli* (**Figure 5A**), retarded *G*. adiacens growth (**Figure 5B**) and did not influence on *Lactobacillus salivarius* and *Lactobacillus crispatus* viability (**Figure 5C and D**) demonstrating that fluconazole effect might be strain and species dependent.

**Figure 4:**
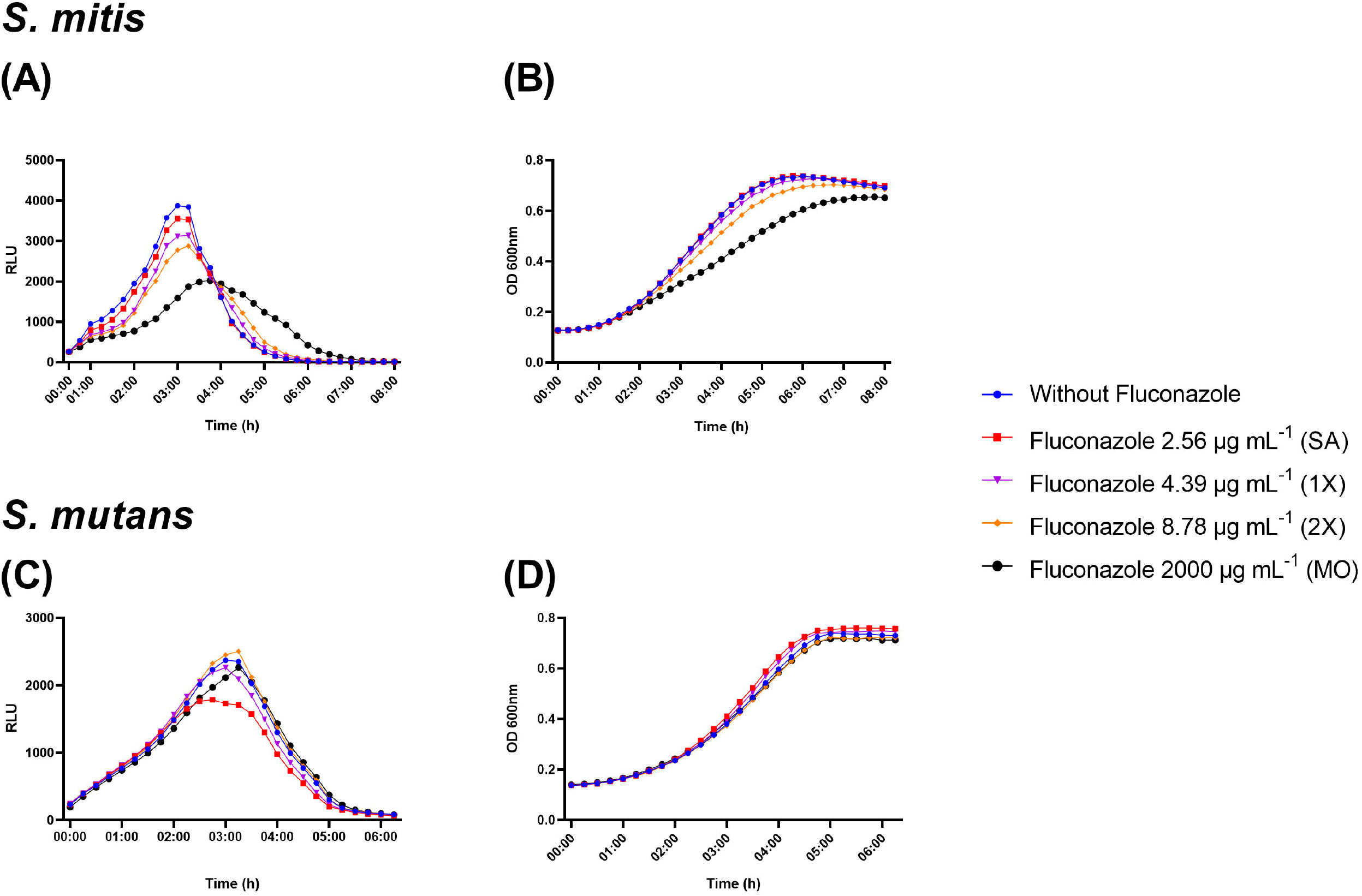
Real-time viability and metabolic activity assay showing *S. mitis* and *S. mutans* growth evaluated as relative light unit (RLU) by the expression of _*p*_*ldh-luc* reporter strains (A and C) and growth by optical density (OD; B and D). Bacterial cells were grown in the presence of fluconazole at 2.56 μg.mL^-1^ (SA), 4.39 μg.mL^-1^ (1x - peak plasma concentration), 8.78 μg.mL^-1^ (2x - twice peak plasma concentration) and 2000 μg.mL^-1^ (MO). Samples without exposure to fluconazole were included as negative control.

**Figure 5:**
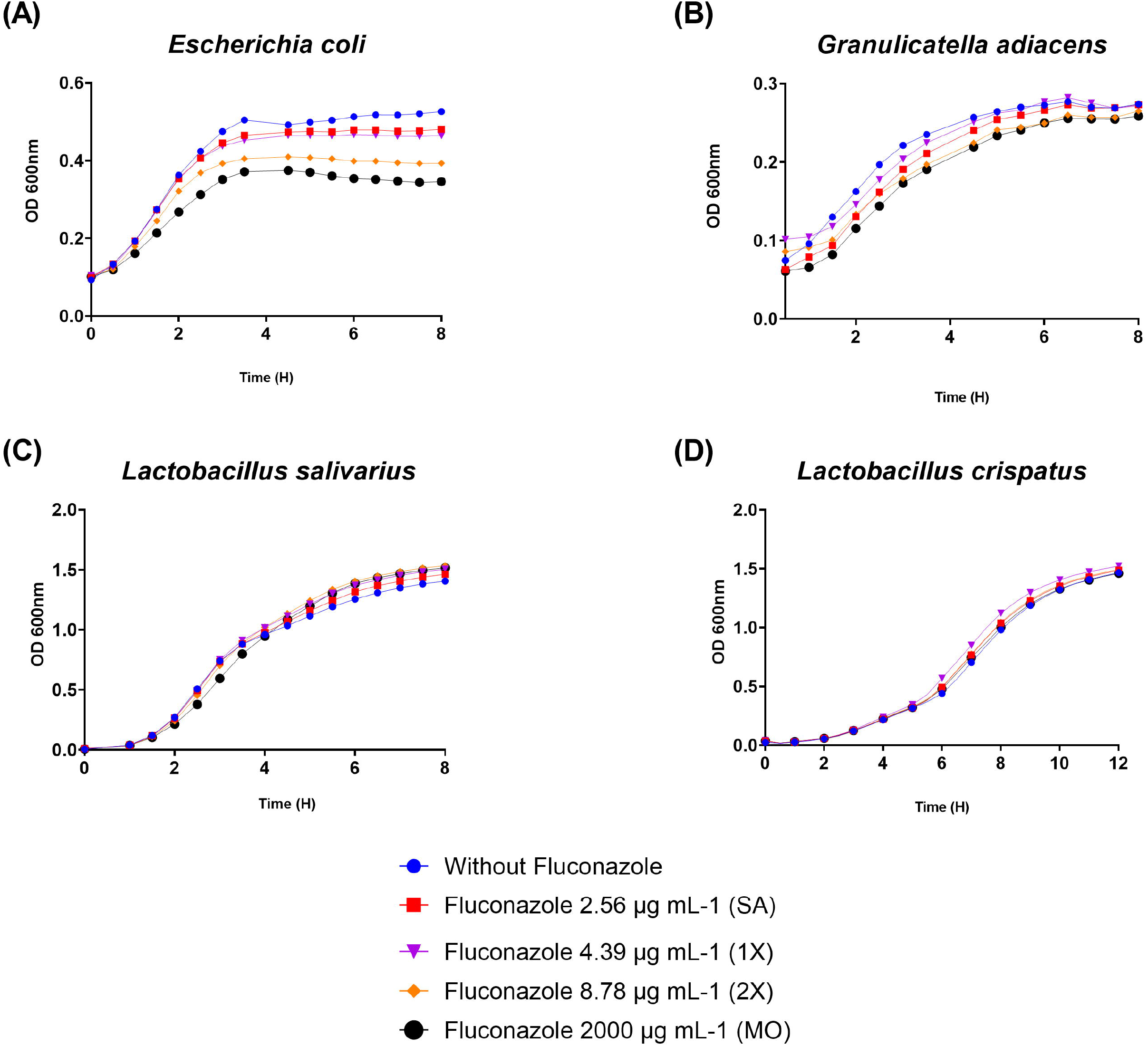
Planktonic growth results assessed by optical density (OD) showed that fluconazole at 2.56 μg.mL^-1^ (SA), 4.39 μg.mL^-1^ (1x), 8.78 μg.mL^-1^ (2x) and 2000 μg.mL^-1^ (MO) reduced planktonic growth of *E. coli* **(Figure 5A)**, retarded *G. adiacens* growth **(Figure 5B)** and did not influence on *Lactobacillus salivarius* and *Lactobacillus crispatus* viability **(Figure 5C,D)**.

### 3.3 Effect of fluconazole on *S. mitis* biofilms

In the *S. mitis* biofilm assay, fluconazole had no significant effect on CFU at any time or concentration (p>0.05) (**Figure 6A**). However, fluconazole significantly altered the biofilm dry biomass by initially reducing it at 6 h, in parallel with a reduced drop in pH, followed by stabilization by 24 h and subsequent increase by 48 h compared to the control samples (**Figure 6B**). In the presence of fluconazole, the pH dropped slower for the 3 highest concentrations compared to the negative control at 6 h. At 24 h the pH for the highest concentration was still higher than the control, while at 48 h no significant differences were observed (**Figure 6C**).

**Figure 6:**
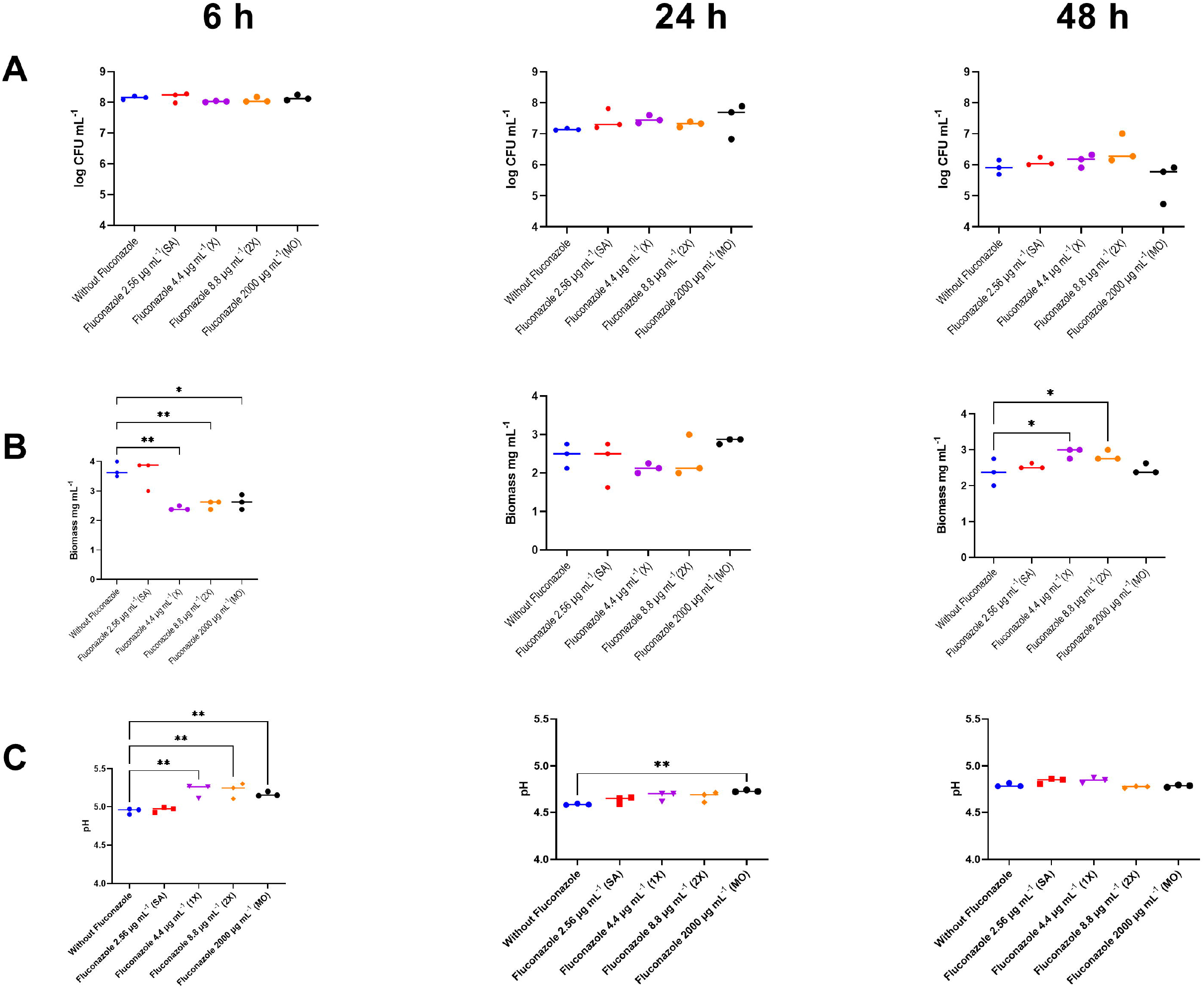
CFU, dry biomass and pH of *S. mitis* biofilms grown for 6, 24, and 48 h in the presence of fluconazole at 2.56 μg.mL^-1^ (SA), 4.39 μg.mL^-1^ (1x), 8.78 μg.mL^-1^ (2x) and 2000 μg.mL^-1^ (MO), or without fluconazole exposure (negative control). Statistical difference *p<0.05, **p<0.01 compared to the control group using One-Way ANOVA followed by Tukey’s multi-comparison post hoc test. 512

## 4 Discussion

Antimicrobials can have unwanted side effects related to their impacts on the host cells or non-targeted microorganisms. By using an *ex-vivo* human microbiota model that reproduces the large species diversity found in oral biofilms (25), we demonstrate fluconazole effects on biomass and viability of oral microbial communities in the absence of host factors. Effects on selected streptococcal and non-streptococcal strains were also observed, highlighting the potential of fluconazole to have direct collateral effects on off-target microbes.

The reduction in biomass and increased CFU levels in the oral microbiota model were not accompanied by changes in pH or total DNA load in the microbiota samples. Although the four factors are inter-linked, each methodology requires specific interpretation. Dry biomass measurements report the total weight of dehydrated biofilms, which is influenced by the amount and type of microbes and also by the extracellular matrix fraction. CFU on the other hand provides relevant information on viable cells able to grow under the specific growth conditions. Total DNA gives in turn an estimate of microbial load, but it does not differentiate between live and dead cells, or between intracellular and extracellular DNA. Overall, the results highlight the complexity of phenotypes influenced by fluconazole exposure. At the microbial community level, changes in the proportion of *V. atypica, V. dispar, Lactobacillus*, and *F. nucleatum* were observed. *In vivo*, a variety of additional factors implicated in microbial community responses are expected to influence microbiota composition, including conditions associated with the use of fluconazole, such as for instance denture-related stomatitis by Candida spp, as well as numerous host and environmental factors. Thus, the model used in our study is not intended to extrapolate the results to *in vivo* conditions, but to highlight that bacterial communities can be affected by fluconazole by mechanisms other than its antifungal activities. To our knowledge, this is the first study to investigate the potential antibacterial impact of fluconazole on complex human-derived microbiomes and on oral species. For the single species experiments, the choice of *S. mitis* was due to its relevance as one of the most prevalent and abundant species in oral microbiomes, and of *S. mutans* due to its symbiotic association with Candida spp. (32). Non-streptococcal species were further evaluated to better understand the specificity of the effects in single species models. Future studies examining the potential impact of fluconazole on other species are warranted.

The most studied class of antimicrobials are antibiotics. Antibiotic stress triggered by sub-inhibitory concentrations has been shown to enhance biofilm formation by different bacteria, including streptococci and other major bacterial human colonizers (33,34). Recent studies indicate that antifungal agents in low concentrations, can have a parallel effect on fungi, by influencing fungal biofilm formation (35). In our model, the concentration of fungal DNA extracted from the microbiota and analyzed by RT-PCR was below the minimal value for reliable detection in (< 20 fg) both for controls and fluconazole treated samples. The finding that bacteria constituted the large majority of the population, led us to hypothesize that the fluconazole effect could be related to a direct off-target effect on bacteria. We then investigated the potential antibacterial effect of fluconazole using *Streptococcus mitis*, the most prevalent and abundant streptococcal colonizer of the oral cavity, and *Streptococcus mutans*, an important pathogen in dental caries.

By combining standard culture density measurements and a luciferase reporter system that enables non-disruptive analyses of the metabolic status and viability of streptococci in real-time (27,31), we found that fluconazole had no or only a minimal effect on *S. mutans* planktonic growth, whereas for *S. mitis* there was a concentration dependent inhibition in growth and in viability/metabolic activity. The effect on metabolic activity was also supported by time measurements showing a delay in pH drop in the supernatants of *S. mitis* biofilms exposed to fluconazole. Additionally, the results of *Escherichia coli, G. adiacens, L. crispatus* and *L. salivarius* planktonic growth demonstrate that the effect of fluconazole could be strain and species dependent. The molecular mechanisms underlying this matter remain unknown and future investigations should address whether such effects extend to other bacterial species and strains.

Like the oral microbiota biofilm model, exposure to fluconazole resulted in a reduction in biofilm mass by *S. mitis*, but only at the initial phase of biofilm formation (6 h). At 48h, most of the samples grown in the presence of fluconazole had a higher biomass than the control. Although a direct comparison between the human microbiota model and the single-species model used in our study is not possible given the inherent methodological differences, the results indicate that fluconazole has the potential to impact both complex- and single species-microbial communities.

Azole antifungals work by altering the fungal cell membrane via potent inhibition of cytochrome P450 lanosterol demethylases in the membranes of fungi. In bacteria, P450 was initially found in the Actinomycetes group, including Mycobacterium tuberculosis, other mycobacteria, and selected Streptomyces species (18). Binding of azoles to a P450 enzyme of Mycobacteria has been demonstrated for fluconazole and several other azoles (18). With the advancements in next generation sequencing in more recent years, it has become apparent that P450 enzymes are more widespread in bacteria than previously realized (36,37). In a large screen within the Firmicutes group, the presence of P450 was found in 24 % of the species (37). However, it remains largely unexplored their functions in bacteria and whether any of these can be targets by azoles. An exception are the selected species in the Actinomycetes group described above, for which strong inhibition has been demonstrated in different studies (18). In streptococci, one of the most prominent colonizers of the oral cavity, P450 is not found in any of the species. Since *S. mitis* apparently lacks P450 enzymes, the fluconazole effects observed in *S. mitis* would involve alternative targets.

Fluconazole’s resistance breakpoint concentrations for *Candida* spp. were recently investigated for their antibacterial effect against selected gram positive and gram-negative species (38). While quaternary ammonium derivatives of fluconazole had a potent inhibitory effect, none of the bacteria tested were inhibited by fluconazole alone. However, the experiments were based on standard methods for MIC determination that examine inhibition only when the bacteria have reached the stationary phase of growth. Another study showed that in plate agar assays, fluconazole inhibited *S. lividans*, although to a lesser extent than imidazoles (18). Our results indicate that although the effects on *S. mitis* were not observed at stationary phase, they were present at earlier growth phases. Accordingly, measurements of density and viability of *S. mitis* revealed a delay in reaching stationary phase in the presence of fluconazole. Such inhibitory effects would have been missed by standard single-time methods to determine MIC values.

Taken together, our results demonstrate that fluconazole at clinically relevant concentrations can selectively impact biofilm and planktonic growth of both complex- and single species-microbial communities, and therefore act as a microbial stressor. Numerous environmental stressors, including antibiotics, have been associated with increased biofilm formation and responses that induce resistant phenotypes in bacteria. The lack of studies examining fluconazole effects during bacterial growth indicates that the effect of fluconazole as a stressor deserves future attention. Further knowledge on such effects will contribute to a better understanding of dysbiosis and resistant phenotypes induced by antimicrobials, with potential implications for the development of superior azole-based inhibitors.

## 6 Author Contributions

Conceptualization, Dornelas Figueira, LM and Petersen, FC; methodology, Dornelas Figueira, LM; Ricomini Filho, AP, Junges, R and Åmdal, HA; resources, Del Bel Cury, AA and Petersen, FC; data curation, Petersen, FC and Ricomini Filho, AP; writing original draft preparation, Dornelas Figueira, LM; writing review and editing, Ricomini Filho, AP and Petersen, FC; supervision, Del Bel Cury, AA and Petersen, FC; project administration, Del Bel Cury, AA and Petersen, FC; funding acquisition, Del Bel Cury, AA, Ricomini Filho, AP and Petersen, FC. All authors have read and agreed to the published version of the manuscript.

## 8 Institutional Review Board Statement

The study was conducted in accordance with the Declaration of Helsinki and approved by the Norwegian Regional Ethics Committee – REK Nord (REK20152491) for studies involving human samples.

## 9 Acknowledgments

This study was financed in part by the **Coordination for the Improvement of Higher Education** - Brazil (CAPES) - Finance Code 001. The authors also thank CAPES for the scholarship provided to the first author (grant number 88887.369710/2019-00 and 88887.508558/2020-00) as well as the **Research Council of Norway and the Norwegian Agency for International Cooperation and Quality Enhancement in Higher Education (DIKU)** for providing financial funds for the development of this study and the first author scholarship co-participation (RESISPART project, grant 274867 and RESISFORCE project, grant 322375). The authors also thank the **Brazilian National Council for Scientific and Technological Development** (CNPq) for the financial support provided to the fifth author (grant number 306275/2016-3). The authors thank Dr. Jacqueline Abranches for providing part of bacterial strains used in this manuscript.

